# Neuroinvasion and encephalitis following intranasal inoculation of SARS-CoV-2 in K18-hACE2 mice

**DOI:** 10.1101/2020.12.14.422714

**Authors:** Pratima Kumari, Hussin A. Rothan, Janhavi P. Natekar, Shannon Stone, Heather Pathak, Philip G. Strate, Komal Arora, Margo A. Brinton, Mukesh Kumar

**Affiliations:** Department of Biology, College of Arts and Sciences, Georgia State University, Atlanta, Georgia 30303

**Keywords:** SARS-COV-2, COVID-19, K-18ACE2 mice, Neuroinvasion, Neuroinflammation, Encephalitis

## Abstract

Severe acute respiratory syndrome coronavirus-2 (SARS-CoV-2) infection can cause neurological disease in humans, but little is known about the pathogenesis of SARS-CoV-2 infection in the central nervous system. Herein, using K18-hACE2 mice, we demonstrate that SARS-CoV-2 neuroinvasion and encephalitis is associated with mortality in these mice. Intranasal infection of K18-hACE2 mice with 10^5^ plaque-forming units of SARS-CoV-2 resulted in 100% mortality by day 6 after infection. The highest virus titers in the lungs were observed at day 3 and declined at days 5 and 6 after infection. In contrast, very high levels of infectious virus were uniformly detected in the brains of all the animals at days 5 and 6. Onset of severe disease in infected mice correlated with peak viral levels in the brain. SARS-CoV-2-infected mice exhibited encephalitis hallmarks characterized by production of cytokines and chemokines, leukocyte infiltration, hemorrhage and neuronal cell death. SARS-CoV-2 was also found to productively infect cells within the nasal turbinate, eye and olfactory bulb, suggesting SARS-CoV-2 entry into the brain by this route after intranasal infection. Our data indicate that direct infection of CNS cells together with the induced inflammatory response in the brain resulted in the severe disease observed in SARS-CoV-2-infected K18-hACE2 mice.

## 1. Introduction

Severe acute respiratory syndrome coronavirus-2 (SARS-CoV-2) infection in humans can cause pneumonia, acute respiratory distress syndrome, acute lung injury, cytokine storm syndrome and death [1,2]. Although SARS-CoV-2 infection primarily causes respiratory disease; some patients develop symptoms of neurological disease, such as headache, loss of taste and smell, ataxia, meningitis, cognitive dysfunction, memory loss, seizures and impaired consciousness [3–10]. SARS-CoV-2 infection also induces long-term neurological sequelae in at least one-third of human cases. Infection with other coronaviruses, such as mouse hepatitis virus (MHV) in mice, and SARS-CoV-1 and Middle East Respiratory Syndrome (MERS) in humans have been shown to cause neurological disease [11,12]. However, little is known about the pathophysiology of SARS-CoV-2-associated neurological disease in humans.

Central nervous system (CNS) cells that express the SARS-CoV-2 receptor, angiotensin-converting enzyme 2 (ACE2), include neurons, glial cells and astrocytes [13,14]. ACE2 is expressed in multiple human brain areas, including the amygdala, cerebral cortex and brainstem with the highest expression levels found in the pons and medulla oblongata in the brainstem that contain the medullary respiratory centers of the brain [8,15]. Several human autopsy reports have documented the presence of SARS-CoV-2 RNA in brain tissues [16,17]. Human iPSC derived neural progenitors cells (NPCs) have been shown to be permissive to SARS-CoV-2 infection and both viral proteins and infectious viral particle production were detected in neurospheres and brain organoids infected with SARS-CoV-2 [18,19]. Human autopsy reports have shown evidence of lymphocytic panencephalitis, meningitis and brainstem perivascular and interstitial inflammatory changes with neuronal loss in COVID-19 patients [20]. These data suggest that SARS-CoV-2 can productively infect human CNS cells [21]. However, the contributions of CNS cell infection and induced neuroinflammation to the pathogenesis of SARS-CoV-2-associated disease are not well understood.

Small animal models provide a means for studying the neurological complications associated with SARS-CoV-2 infection. It was recently reported that intranasal inoculation with SARS-CoV-2 results in a rapidly fatal disease in K18-hACE2 mice [22–25]. These studies were focused on describing the acute lung injury in SARS-CoV-2 infected K18-hACE2 mice that was associated with high levels of inflammatory cytokines and accumulation of immune cells in the lungs [22–25]. In these published studies, infectious virus or viral RNA was not detected in the olfactory bulbs or brains of the majority of the infected animals, indicating restricted neurotropism of SARS-CoV-2 in K18-hACE2 mice. In the present study, we show that intranasal infection of six-week-old K18-hACE2 mice by SARS-CoV-2 can cause severe neurological disease with the brain being a major target organ for infection by this route of infection and neuroinflammation and neuronal death contributing to the infection-associated morbidity and mortality. The data also suggest that the SARS-CoV-2 can be trafficked to the brain via the olfactory bulb with subsequent transneuronal spread, as has been reported for other coronaviruses [26,27].

## 2. Materials and Methods

### Mice

Hemizygous K-18 hACE2 mice were purchased from the Jackson Laboratory (Bar Harbor, ME). All the animal experiments were conducted in a certified animal biosafety level-3 (ABSL-3) laboratory at the Georgia State University (GSU). The protocol was approved by the GSU IACUC (Protocol number A20044). Six-week-old hemizygous K-18 hACE2 mice were infected with 10^5^ plaque-forming units (PFU) of SARS-CoV-2 strain USA-WA1/2020 under ABSL-3 containment by intranasal inoculation. SARS-CoV-2 (USA-WA1/2020) was isolated from an oropharyngeal swab from a patient in Washington, USA (BEI NR-52281) [28]. Animals in the control group received equivalent amounts of sterile PBS via the same route. Roughly equal numbers of male and female mice were used. Animals were weighed and their appetite, activity, breathing and neurological signs assessed twice daily [29,30]. Mice that met the human endpoint criteria were euthanized to limit suffering. In independent experiments, mice were inoculated with PBS (Mock) or SARS-CoV-2 intranasally, and on days 1, 3, 5 and 6 after infection, animals were anesthetized using isoflurane, perfused with cold PBS and respiratory (nasal turbinate and lung) and other tissues (spleen, heart, liver, kidney, pancreas, eye, olfactory bulb and brain) were collected and flash frozen in 2-methylbutane (Sigma, St. Louis, Missouri, United States) [31–33]. Alternatively, mice were perfused with PBS followed by 4% paraformaldehyde (PFA) and tissues were harvested, cryoprotected in 30% sucrose (Sigma, St. Louis, Missouri, United States), and embedded in optimum cutting temperature (OCT) as described previously [31,34].

### Quantification of the virus load

The virus titers were analyzed in the tissues by plaque assay and quantitative real-time PCR (qRT-PCR) [28,29]. Briefly, frozen tissues were weighed and homogenized in a bullet blender (Next Advance, Averill Park, New York, United States) using glass or zirconium oxide beads. Virus titers in tissue homogenates were measured by plaque assay using Vero cells. Quantitative RT-PCR was used to measure viral RNA levels using primers and probes specific for the SARS-CoV-2 N gene as described previously [28]. Viral genome copies were determined by comparison to a standard curve generated using a known amount of RNA extracted from previously titrated SARS-CoV-2 samples. Frozen tissues harvested from mock and infected animals were weighed and lysed in RLT buffer (Qiagen) and RNA was extracted using a Qiagen RNeasy Mini kit (Qiagen, Germantown, MD, USA). Total RNA extracted from the tissues was quantified, normalized and viral RNA levels per µg of total RNA were calculated.

### Measurement of cytokines, chemokines and interferons

The levels of mRNA for select cytokines/chemokines (IL-1β, IL-6, TNF-α, IFN-γ, CCL2 and CCL3) and interferon-α (IFN-α) were determined in total RNA extracted from the lungs and brain using qRT-PCR. The fold-change in infected tissues compared to mock tissues was calculated after normalizing to the GAPDH gene [31,34]. The primer sequences and annealing temperatures used for qRT-PCR are listed in Table 1. The protein levels of IFN-α were measured in the lung and brain homogenates using an ELISA kit (PBL Interferon Source, Piscataway, NJ, USA) [29,30].

**Table 1.**
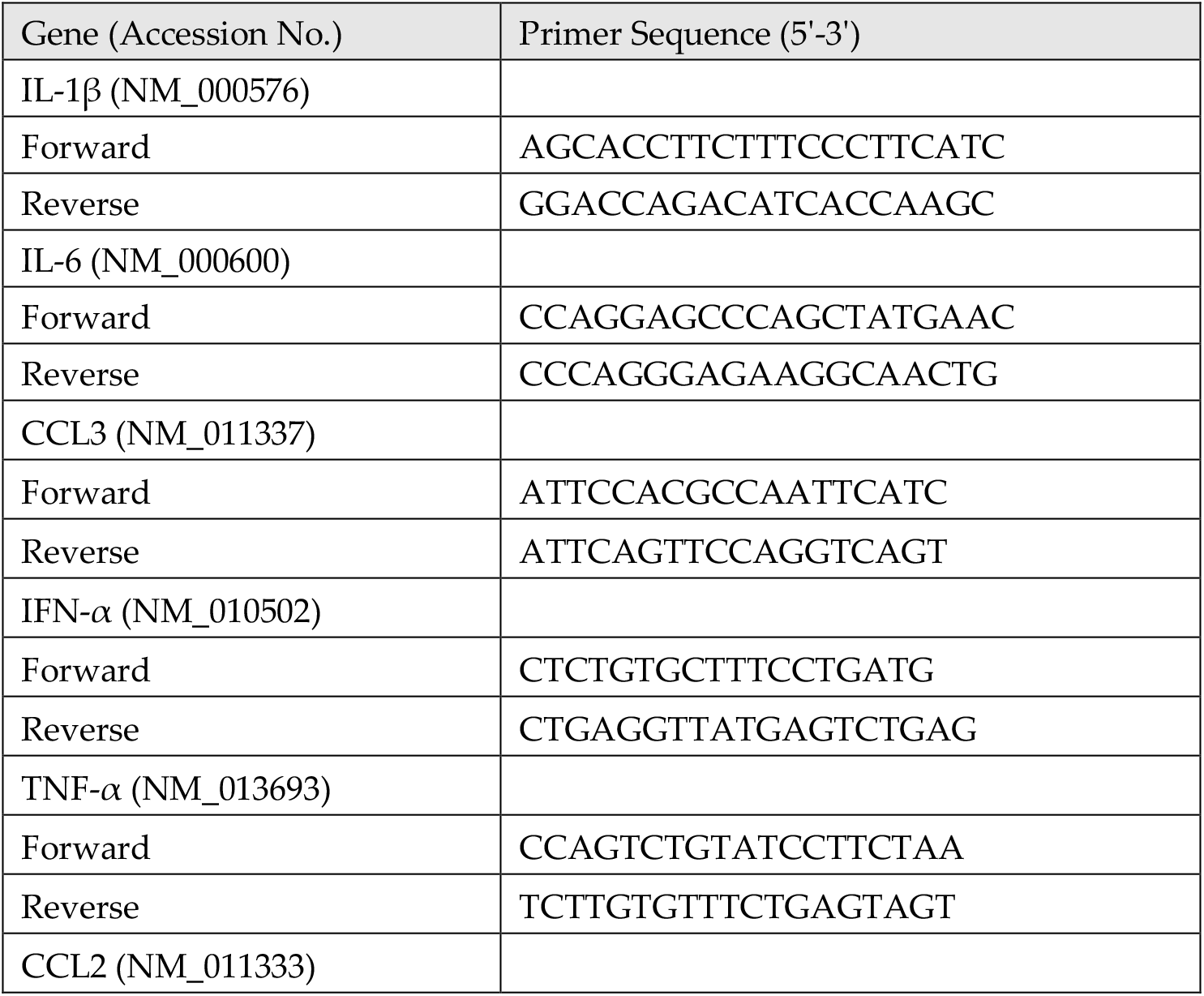
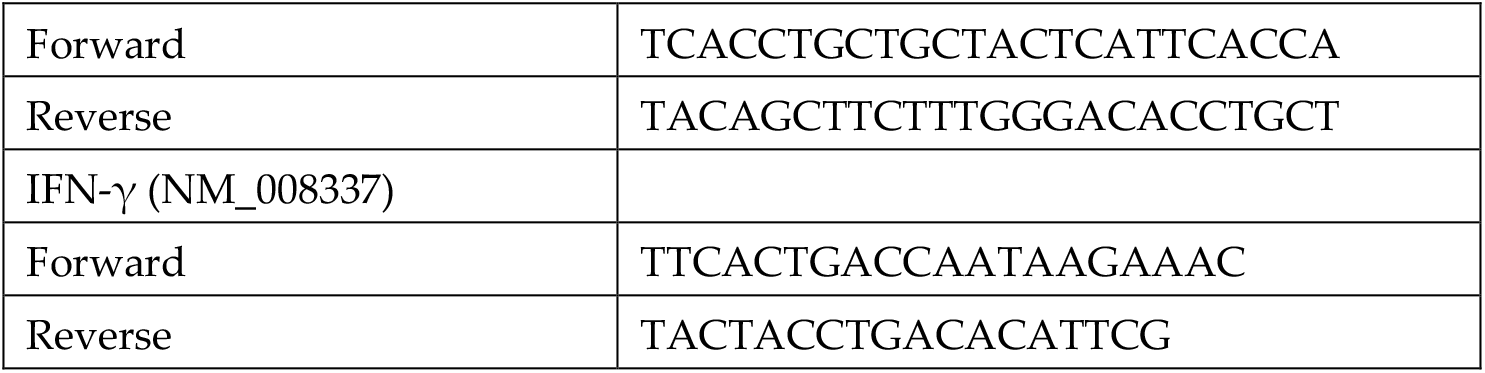
Primer sequences used for qRT-PCR.

### Immunohistochemistry

Sagittal sections (10-µm thick) were cut from the hemi-brain tissues frozen in OCT. Tissue sections were stained with hematoxylin and eosin (H&E) for histopathological evaluation [34,35]. Additionally, tissue sections were incubated with anti-CD45, anti-NeuN and anti-SARS-CoV-2 spike protein antibodies (Thermo Fisher Scientific, Norcross, GA, USA) overnight at 4°C followed by incubation with Alexa Fluor 546- or Alexa Fluor 488-conjugated secondary antibody for 1 hr at room temperature [31,34]. Terminal deoxynucleotidyl transferase dUTP nick end labeling (TUNEL) staining was conducted using an in-situ cell death detection kit (Roche, Indianapolis, Indiana, United States) as per the manufacturer’s instructions [31,34]. Images were acquired using the Invitrogen™ EVOS™ M5000 Cell Imaging System (Thermo Fisher Scientific, Norcross, GA, USA).

### Statistical Analysis

An unpaired Student’s t-test was used to calculate p values of significance. Differences with P values of <0.05 were considered significant.

## 3. Results

### Characteristics of K18-hACE2 mice following SARS-CoV-2 infection by the intranasal route

Six-week-old K18-hACE2 mice of both sexes were infected intranasally with PBS (mock, n=10 mice) or 10^5^ PFU of SARS-CoV-2 in PBS (n=20 mice). The mock-infected mice remained healthy throughout the observation period. Virus infection resulted in 100% mortality by day 6 after infection (Figure 1A). Infected mice experienced significant weight loss on days 4 through 6 after infection compared to the mock-infected group (Figure 1B). Starting on day 4, all the infected animals began to show signs of disease, such as lethargy, slow movement and labored breathing. Neurological symptoms, such as hunchbacked posture, ruffled fur, tremors and ataxic gait, were also observed in infected mice on days 5 and 6 after infection.

**Figure 1.**
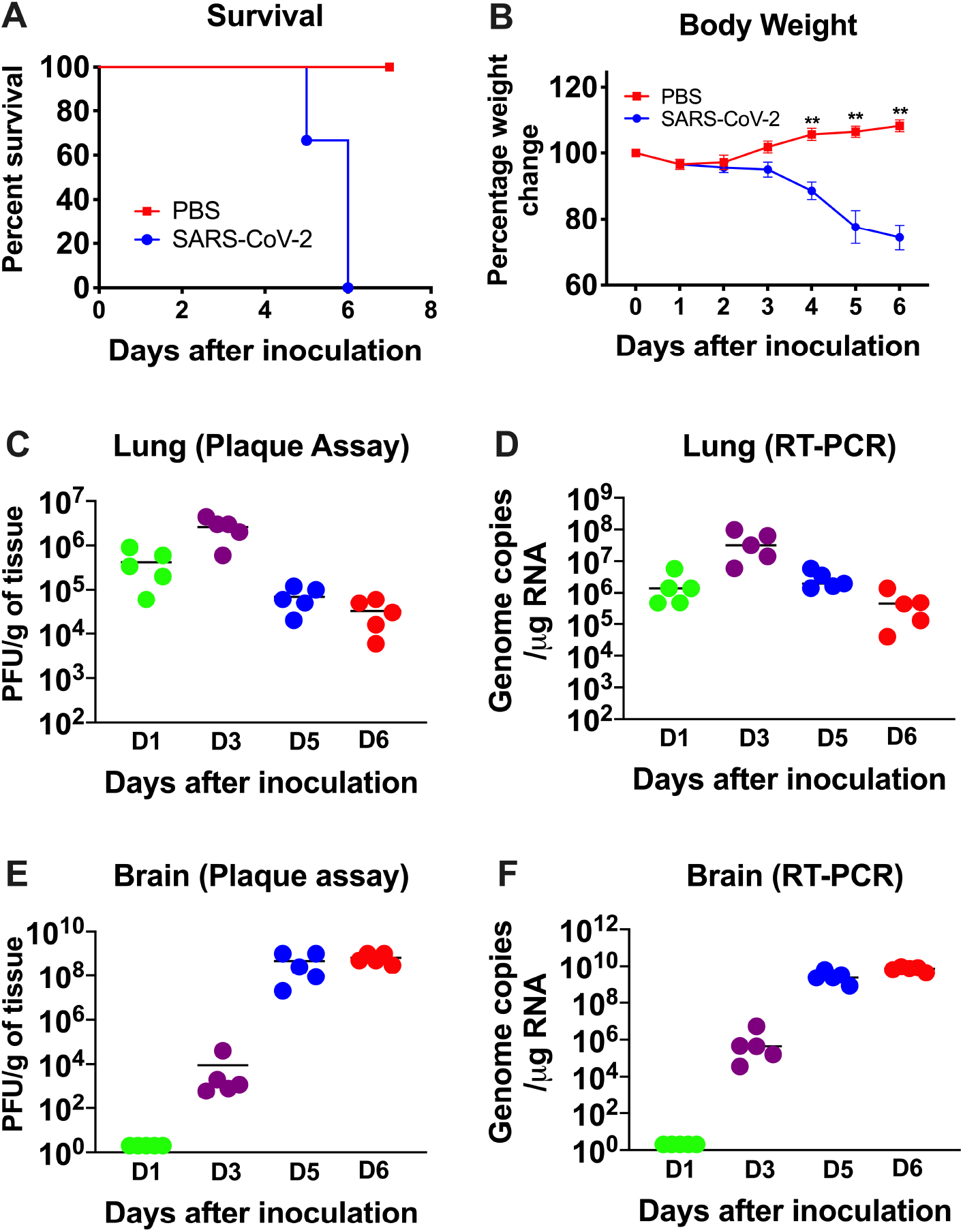
Analysis of survival, body weight and virus titers in K18-hACE2 mice following SARS-CoV-2 infection. K18-hACE2 mice were inoculated intranasally with SARS-CoV-2 (10^5^ PFU, n=20) or PBS (Mock, n=10). (A) Percent survival was determined. (B) Percent daily body weight change in the animals. Error bars represent SEM. **p < 0.001. The kinetics and levels of SARS-CoV-2 were determined in the lungs (C and D) and brain (E and F) by plaque assay and qRT-PCR. The data are expressed as PFU/g of tissue or genome copies/µg of RNA. Each data point represents an individual mouse. The solid horizontal lines signify the median.

### Virus replication in the periphery and brain of K18-hACE2 mice

Six-week-old K18-hACE2 mice of both sexes were infected intranasally with PBS (mock, n=12 mice) or 10^5^ PFU of SARS-CoV-2 in PBS (n=20 mice) and groups of 5 mice were used to measure the viral loads in the peripheral organs and brain at early (day 1), middle (day 3) and late (days 5 and 6) stages of infection. High virus levels were observed in the lungs on day 1, reached peak levels at day 3 and declined at days 5 and 6 after infection (Figures 1C and 1D). In contrast, virus was not detected in the brain on day one but was present by day 3 after infection. Very high levels of viral RNA and infectious virus were detected in the brains of all the animals by days 5 and 6 after infection (Figures 1E and 1F). The onset of neurological symptoms and mortality in infected mice correlated with peak virus titers in the brain.

The virus replication kinetics observed in the nasal turbinates was similar to those in the lungs with the highest viral RNA levels detected during the early stage of infection (days 1 and 3), followed by a decline at later stages of infection (days 5 and 6) (Figure 2). In the olfactory bulbs and eyes, low levels of viral RNA were detected on days 1 and 3 after infection, with very high levels of viral RNA detected in all the animals on days 5 and 6 after infection indicating productive infection within the olfactory system (Figure 2). In contrast, little if any virus was detected in the serum of infected mice at any time after infection tested.

**Figure 2.**
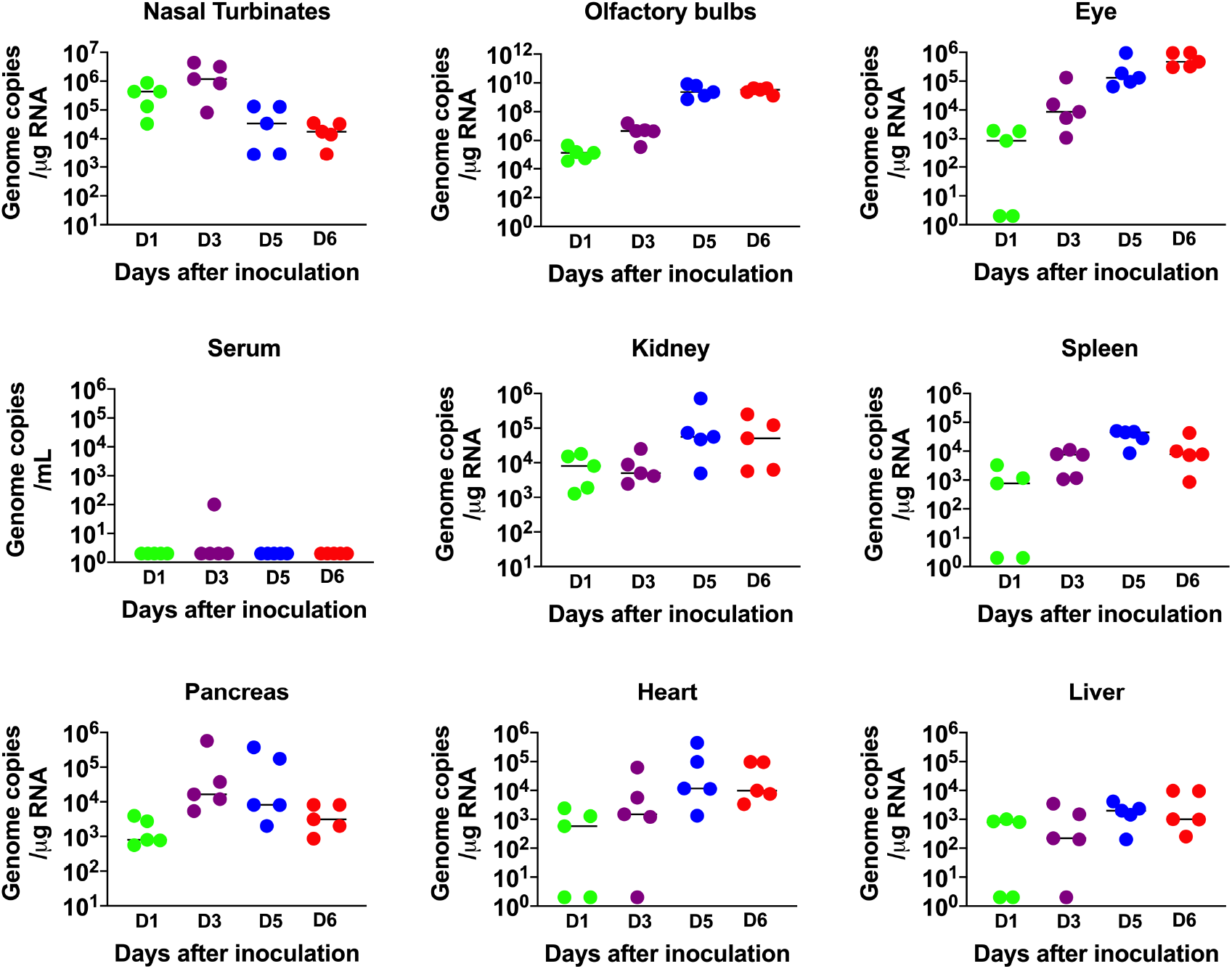
Analysis of virus tropism in K18-hACE2 mice. The viral RNA copy number in the nasal turbinates, olfactory bulbs, eye, serum, kidney, spleen, pancreas, heart and liver was determined on days 1, 3, 5 and 6 after infection by qRT-PCR and expressed as genome copies/µg of RNA. Each data point represents an individual mouse. The solid horizontal lines signify the median.

Since the K18 promoter is known to be active in the epithelium of multiple organs of K-18-hACE2 mice [27,36], we also evaluated viral RNA levels in other peripheral organs. Viral RNA was detected in the heart, kidney, spleen, pancreas and liver on days 1 and 3. There was a slight increase in RNA levels on days 3 and 5 in each of these organs, suggesting limited virus replication at these sites (Figure 2). These data agree with previous reports that also showed the presence of SARS-CoV-2 RNA in these organs [22-25,27,36].

### Inflammatory changes in the lungs and brain of SARS-CoV-2-infected mice

IFN signaling has a pivotal role in developing an innate and adaptive immune response to viral infection [37,38]. Therefore, we measured the mRNA and protein levels of IFN-α in the lungs and brain. In the lungs, an increase in both IFN-α mRNA and protein levels was detected on day 1, peaked at day 3 and then decreased on day 6 after infection (Figures 3A and 3B). In contrast, an IFN response was not detected in the brain on days 1 and 3 after infection. High levels of IFN-α were detected in the brain only by days 5 and 6 after infection (Figures 3C and 3D). Overall, the induction of IFN-α correlated with the SARS-CoV-2 replication kinetics in the lungs and brain. It is interesting to note that relative IFN-α levels were comparatively higher in the lungs compared to the brains of the infected animals despite higher virus replication in the brain.

**Figure 3.**
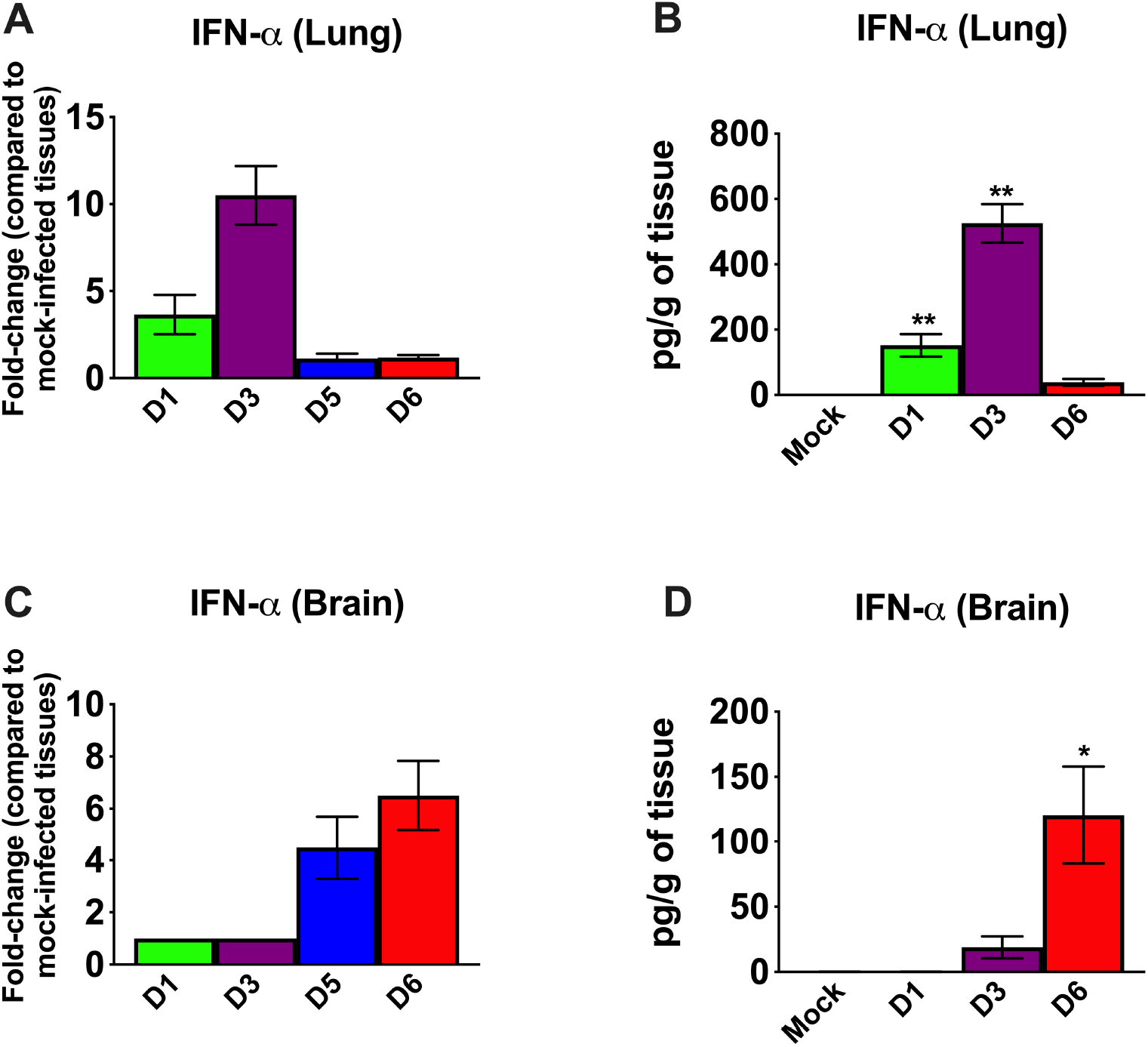
Analysis of mRNA and protein levels of IFN-α in the lungs and brain. The mRNA levels of IFN-α were measured in the lungs (A) and brain (C) by qRT-PCR, and the fold change in the infected tissues compared to the corresponding mock-infected controls was calculated after normalizing to the GAPDH gene. The protein levels of IFN-α were measured in the lungs (B) and brain (D) homogenates using ELISA and expressed as pg/g of tissue. Error bars represent SEM (n = 5 mice per group). *p < 0.05; **p < 0.001.

We next examined the mRNA levels of proinflammatory cytokines and chemokines in the lungs and brain of infected mice. SARS-CoV-2 infection resulted in a 10-fold increase on day 1 and a 100-fold increase on day 3 in the IL-6 mRNA expression in the lungs (Figure 4A). The levels of TNF-α mRNA were elevated ∼10-fold in the lungs on day 1 and 3 after infection. The level of IFN-γ mRNA was elevated by 15-fold on day 3. However, the levels of these cytokines had decreased by day 5 after infection. The IL-1β mRNA levels showed no significant increase at any time point after infection. There was a 100-fold increase in the expression of CCL2 on day 3 (Figure 4B). However, the levels of CCL2 mRNA had decreased by day 5 after infection. CCL3 mRNA levels increased slightly on day 1 and were undetectable at days 5 and 6 after infection.

**Figure 4.**
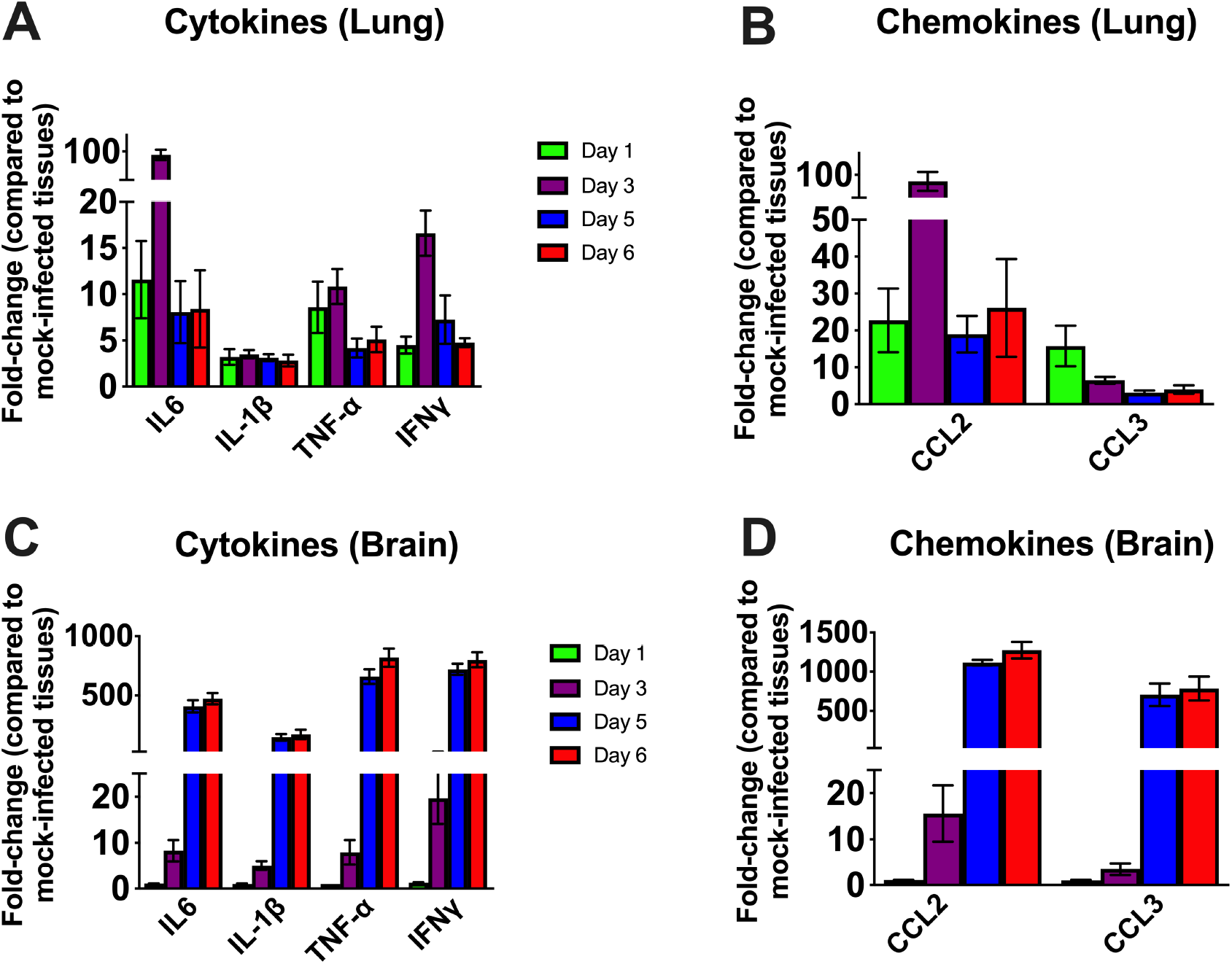
Cytokine and chemokine mRNA level in the lungs and brain. The mRNA levels of various cytokine and chemokine genes were determined in the lungs (A and B) and brain (C and D) using qRT-PCR. Fold change in the infected tissues compared to the corresponding mock controls was calculated after normalizing to GAPDH mRNA in each sample. Error bars represent SEM (n = 5).

In the brain, no increase in the mRNAs of the cytokines or chemokines tested was observed on day 1 after infection. Less than a 10-fold increase was observed in the cytokine mRNA levels on day 3 (Figures 4C). There was a 500-fold increase in IL-6 mRNA by day 5 after infection. TNF-α and IFN-γ mRNA levels increased by ∼ 750-fold in the brain by day 5 after infection. Similarly, IL-1β mRNA levels increased by 400-fold by day 5 after infection. Both CCL2 and CCL3 mRNA levels were elevated by almost 1,000-fold on days 5 and 6 after infection and consistent with the high level of virus in the brain (Figure 4D). These results indicate that the inflammatory response was more pronounced in the brain than in the lungs at the later stage of infection.

### SARS-CoV-2 induced neuropathology in K18-hACE2 mice

We next analyzed the brain sections from infected mice for antigen distribution, infiltration of immune cells and cell death. Immunohistochemical staining for the SARS-CoV-2 spike protein detected cell-associated viral antigen throughout the brain at day 6 after infection. Representative data for sections from the cortex, cerebellum and hippocampus regions are shown in figure 5. We also detected virus antigen in sections of the olfactory bulb of infected animals on day 6. H&E staining of brain sections from the infected mice demonstrated perivascular hemorrhage and neuronal cell death (Figures 6A and 6B). The neurons of infected mice demonstrated shrunken neuron body with light pink cytoplasmic staining representing degenerating neurons (Figure 6B). Enhanced leukocyte infiltration was detected within blood vessel walls and in the perivascular space (Figure 6A). Evidence of leukocyte infiltration was confirmed by direct immunohistochemical analysis of the CD45 antigen, which revealed many CD45-positive cells in the brain parenchyma near neurons (Figure 6C). SARS-CoV-2-induced cell death was evaluated by direct TUNEL staining of brain tissues. On day 6, infected K18-hACE2 mice had elevated numbers of TUNEL-positive cells in the cortex, hippocampus and cerebellum regions, indicating increased cell death (Figure 6D).

**Figure 5.**
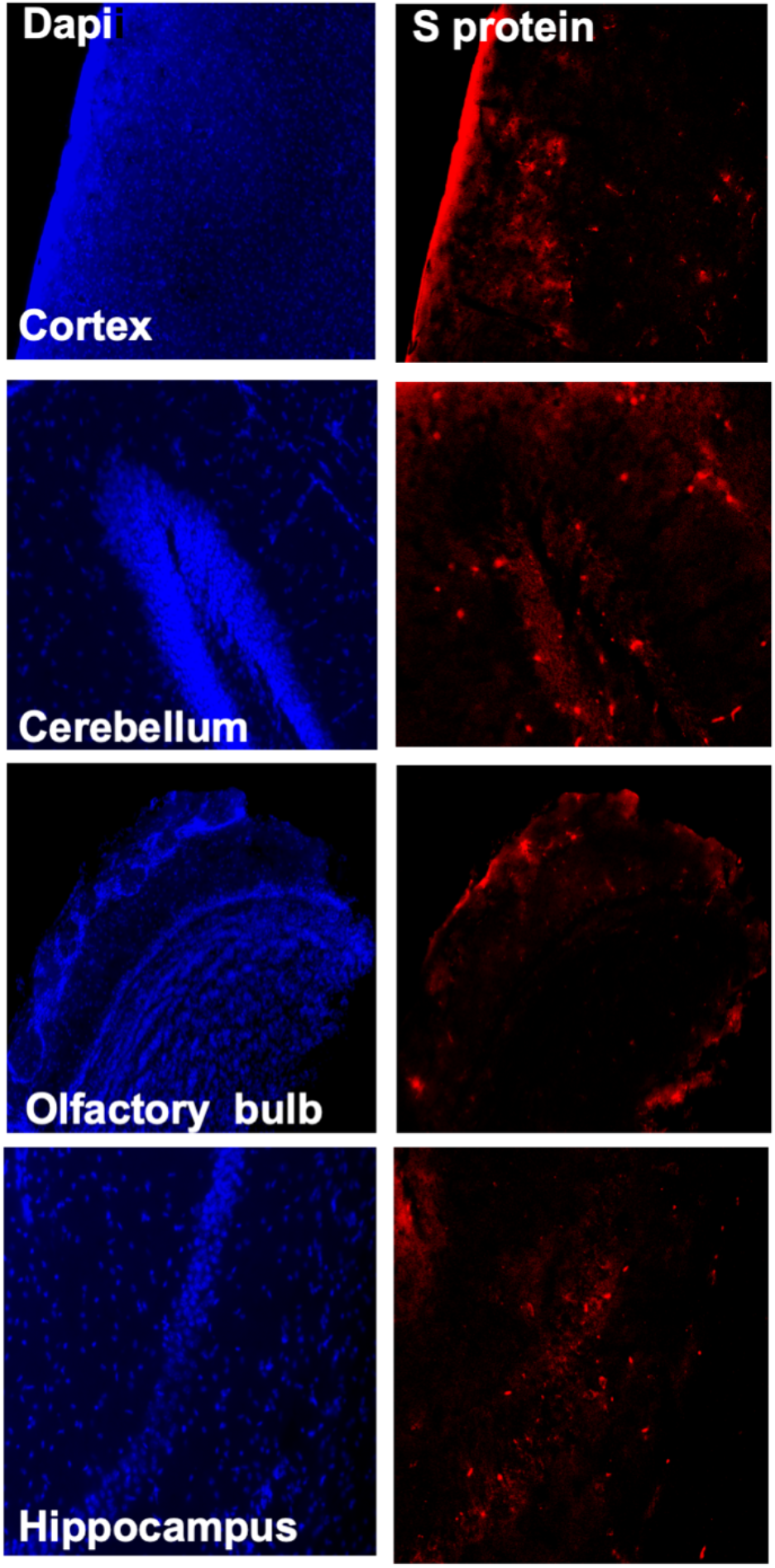
Detection of SARS-CoV-2-infected cells in the brains of K18-hACE2 transgenic mice. Brain sections (day 6 after infection) were stained for SARS-CoV-2 spike protein. Representative immunostaining images showing the presence of SARS-CoV-2 spike protein (red) in the cortex, cerebellum, hippocampus and olfactory bulb of infected mice. Nuclei are stained with DAPI (blue). The photomicrographs shown are representative of the images obtained from five animals. Bars, 20 µm.

**Figure 6.**
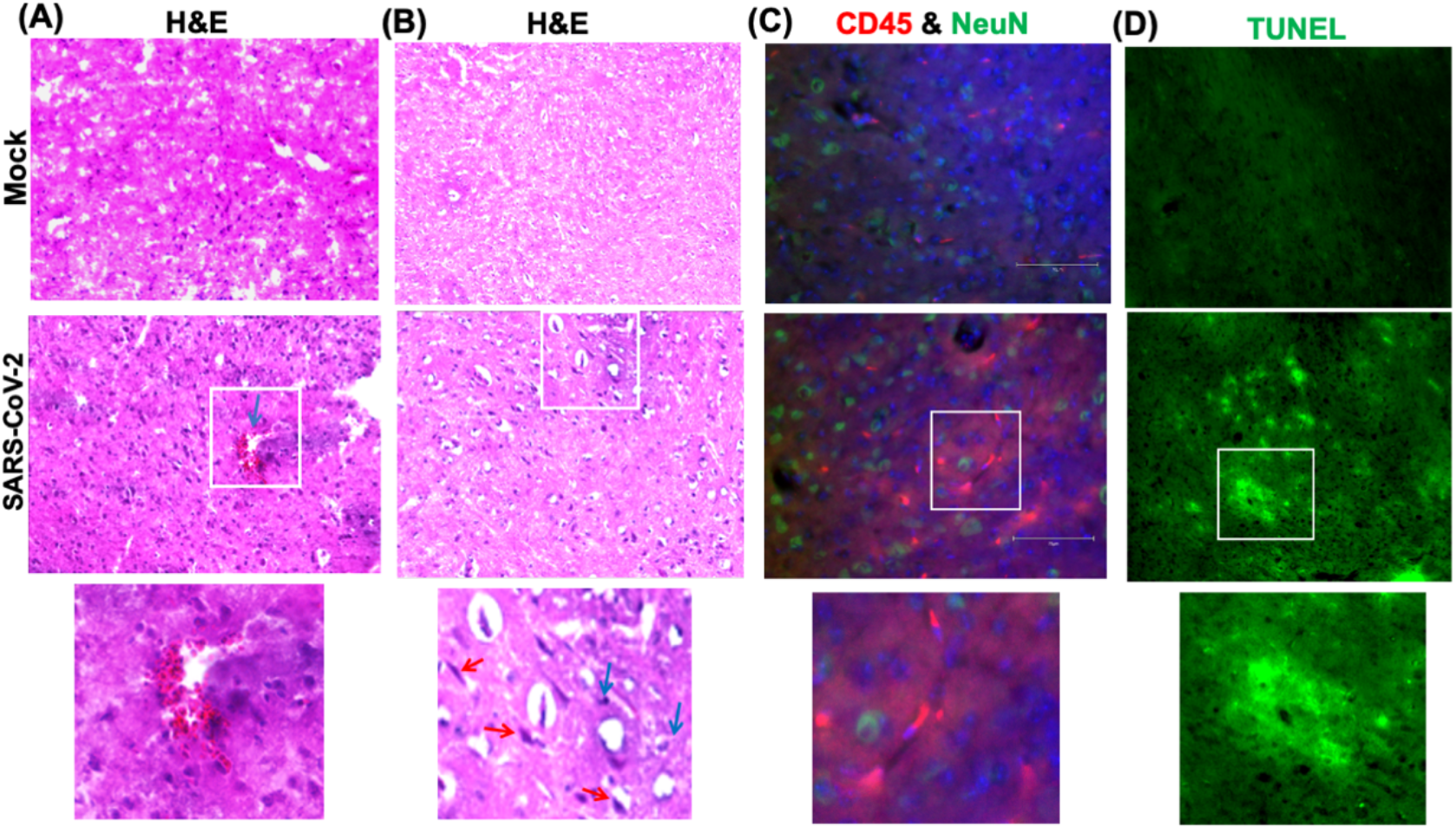
Histopathological analysis of SARS-CoV-2-infected brains. H&E staining of brain sections from mock and SARS-CoV-2-infected mice at day 6 after infection. (A and B) Brain sections show perivascular hemorrhage, enhanced leukocyte infiltration (blue arrows) and neuronal cell death (red arrows). (C) Brain sections were stained for CD45 (Red, leukocyte marker) and NeuN (Green, neuronal cell marker). Nuclei are stained with DAPI (blue). (D) A TUNEL assay was conducted on brain sections from mock and SARS-CoV-2-infected mice at day 6 after infection to detect apoptotic cells. The boxed areas in the second row of panels are enlarged in the bottom row of panels. The photomicrographs shown are representative of the images obtained from five animals. Bars, 20 µm.

## 4. Discussion

This study demonstrates a critical role of direct infection of CNS cells and of the inflammatory response in mediating SARS-CoV-2-induced lethal disease in K18-hACE2 mice. Intranasal inoculation of the virus results in a lethal disease with high levels of virus replication in the brain. Virus infection of the CNS was accompanied by an inflammatory response as indicated by the production of cytokines/chemokines, infiltration of leukocytes into the perivascular space and parenchyma and CNS cell death. Our data also indicate that following infection by the intranasal route, the virus enters the brain by traversing the cribriform plate and infecting neuronal processes located near the site of intranasal inoculation.

Some animal coronaviruses, such as MHV readily infect the neurons and cause lethal encephalitis in mice [11,39]. SARS-CoV infection also induces severe neurological disease after intranasal administration in K18-hACE2 mice [27]. Similarly, in our study, SARS-CoV-2 virus antigen was detected throughout the brain, including the cortex, cerebellum and hippocampus. The onset of severe disease in SARS-CoV-2 infected mice correlated with peak viral levels in the brain and immune cell infiltration and CNS cell death. Peak virus titers in the brains were approximately 1,000 times higher than the peak titers in the lungs, suggesting a high replicative potential of SARS-CoV-2 in the brain. The relative up-regulation of cytokine and chemokine mRNAs was approximately 10 to 50 times higher in the brain compared to the lungs, strongly suggesting that extensive neuroinflammation contributed to clinical disease in mice.

It was recently reported that SARS-CoV-2 infection of K18-hACE2 mice causes severe pulmonary disease with high virus levels detected in the lungs of these mice and that mortality was due to the lung infection [22–25]. In these studies, viral RNA was undetectable in the brains of the majority of the infected animals, indicating a limited role of brain infection in disease induction. An important distinction between our study and others is that we detected high infectious virus titers in the olfactory system and brains of 100% of the infected K18-hACE2 mice. This phenotype was not consistently observed in the aforementioned K18-hACE2 mouse studies [22–25]. Moreover, none of the published studies evaluated the extent of neuroinflammation and neuropathology at the later stages of infection. Our results showed that the inflammatory response was more pronounced in the brain than in the lungs on days 5 and 6 after infection. Although both our study and the previous studies infected mice via the intranasal route, the other studies used older (7- to 9-week-old) K18-hACE2 and a lower viral dose (10^4^ PFU). In our study, six-week-old K18-hACE2 mice were infected with 10^5^ PFU. It appears that initial virus dose and age of animals are critical determinants of tissue tropism in this model. More studies are needed to clarify the parameters that differentially affect tissue tropism, routes of virus dissemination, and mechanisms of lung and brain injuries in K18-hACE2 mice following SARS-CoV-2 infection.

Alterations in smell and taste are features of COVID-19 disease in humans [8,40]. Pathological analyses of human COVID-19 autopsy tissues detected the presence of SARS-CoV-2 proteins in endothelial cells within the olfactory bulb [40,41]. Our data indicate that SARS-CoV-2 can productively infect cells within the nasal turbinate, eye and olfactory bulb in intranasally infected K18-hACE2 mice. Virus infection of cells in these tissues in humans may explain the loss of smell associated with some COVID-19 cases [40]. The detection of virus replication in these tissues suggests that SARS-CoV-2 can access the brain by first infecting the olfactory bulb and then spreading into the brain by infecting connecting brain neuron axons. This hypothesis is consistent with previously published reports that neurotropic coronaviruses infect olfactory neurons and are transmitted to the brain via axonal transportation [8,26,27,42]. Many viruses, such as HSV-1, Nipah virus, rabies virus, Hendra virus and influenza A virus, have also been shown to enter the CNS via olfactory sensory neurons [43–46]. Another route by which a virus can gain access to the brain is via the disruption of the blood-brain barrier (BBB). However, we could not detect any virus in the serum of the infected mice at any time after infection tested, suggesting a limited role of BBB disruption in SARS-CoV-2 neuroinvasion. This finding is in agreement with previously published studies that detected little or no virus in the blood of K18-hACE2 mice after infection with SARS-CoV-1 or SARS-CoV-2 [22-25,27,36].

**In summary**, we found that intranasal Infection of K18-hACE2 mice by SARS-CoV-2 causes severe neurological disease. Our data demonstrate that the CNS is the major target of SARS-CoV-2 infection in K18-hACE2 mice under the conditions used, and that brain infection leads to immune cell infiltration, inflammation and cell death.

## Author Contributions

Conceptualization, M.K., M.A.B; methodology, P.K., J.P.N., H.A.R., S.S., K.A., P.G.S., H.P., M.K.; validation, P.K., J.P.N., H.A.R., S.S., M.K.; formal analysis, P.K., J.P.N., H.A.R., S.S., K.A., M.A.B., M.K.; resources, M.K.; writing—original draft preparation, H.A.R., K.A., M.K.; writing—review and editing, H.A.R., K.A., M.A.B, M.K.; funding acquisition, M.K., M.A.B. All authors have read and agreed to the published version of the manuscript.

## Acknowledgments

We thank members of the GSU High Containment Core and the Department for Animal Research for assistance with the experiments.

